# Representational Connectivity Analysis: Identifying Networks of Shared Changes in Representational Strength through Jackknife Resampling

**DOI:** 10.1101/2020.05.28.103077

**Authors:** Marc N. Coutanche, Essang Akpan, Rae R. Buckser

**Author notes:** Authors contributed equally (Essang Akpan, Rae R. Buckser). Corresponding author: Marc N. Coutanche.

## Abstract

The structure of information in the brain is crucial to cognitive function. The representational space of a brain region can be identified through Representational Similarity Analysis (RSA) applied to functional magnetic resonance imaging (fMRI) data. In its classic form, RSA collapses the time-series of each condition, eliminating fluctuations in similarity over time. We propose a method for identifying representational connectivity (RC) networks, which share fluctuations in representational strength, in an analogous manner to functional connectivity (FC), which tracks fluctuations in BOLD signal, and informational connectivity, which tracks fluctuations in pattern discriminability. We utilize jackknife resampling, a statistical technique in which observations are removed in turn to determine their influence. We applied the jackknife technique to an existing fMRI dataset collected as participants viewed videos of animals (Nastase et al., 2017). We used ventral temporal cortex (VT) as a seed region, and compared the resulting network to a second-order RSA, in which brain regions’ representational spaces are compared, and to the network identified through FC. The novel representational connectivity analysis identified a network comprising regions associated with lower-level visual processing, spatial cognition, perceptual-motor integration, and visual attention, indicating that these regions shared fluctuations in representational similarity strength with VT. RC, second-order RSA and FC identified areas unique to each method, indicating that analyzing shared fluctuations in the strength of representational similarity reveals previously undetectable networks of regions. The RC analysis thus offers a new way to understand representational similarity at the network level.

## 1. Introduction

The principles underlying how information is organized in the brain are central to understanding the neural basis of perceptual, cognitive, and affective states. Through functional magnetic resonance imaging (fMRI), brain activity can be analyzed to understand the representational structure of information within a region, particularly with multivariate analyses, which can probe stimuli and conditions across multiple dimensions (Coutanche, 2013). One such technique, Representational Similarity Analysis (RSA), measures the degree of (dis)similarity between patterns of activity for different stimuli, in order to determine the representational space within a region (Kriegeskorte, Mur, & Bandettini, 2008). This can be represented as a representational dissimilarity matrix (RDM), in which the dissimilarity of activity patterns between stimuli or conditions is reflected.

A common method for comparing the representational spaces of different regions is to measure the similarity (correlation) of RDMs of different regions, through a “second-order RSA” (Kriegeskorte, Mur, & Bandettini, 2008). When comparing RDMs across regions, a second-order RSA compares the overall RSAs (collapsed across time) of different regions (Kriegeskorte, Mur, & Bandettini, 2008). However, combining all timepoints in this manner removes information that may be contained in fluctuations in the strength of representational similarity over time. This is analogous to how the generalized linear model (GLM) examines an average BOLD response. In the case of GLM-based analyses, researchers can also analyze BOLD responses over time to calculate functional connectivity (FC) between regions. FC draws on time-domain information to characterize the interactions between regions within functional networks by identifying shared fluctuations in BOLD activity (Friston, 1997). We have previously shown that multivariate methods can also benefit from analysis of the time domain: fluctuations in multivariate information, such as classifier performance, can be used to identify *informational networks*. Coutanche and Thompson-Schill (2013) illustrated that Informational Connectivity (IC), a method for identifying synchronized fluctuations in the discriminability of multi-voxel patterns over time, identified informational networks that were overlapping but also distinct from the functional networks defined by FC. As in the univariate case, an IC network analysis provides information above and beyond what is available from MVPA alone. Can an equivalent approach be taken for RSA? One key challenge is that collapsing the timepoints must occur to create an RDM; RDMs (by definition) rely on similarity between a given condition and all others, so there is no clear way to extract the RDM for a single time-point alone.

In this paper, we present a novel method to test whether regions are linked through shared fluctuations in the strength of their representational similarity over time. We utilize jackknife resampling, a statistical technique that removes observations one at a time in order to quantify each observation’s effect on the overall (all observations) RDM. The resulting vector represents the influence of each timepoint on a region’s RDM, which may then be compared via correlation to corresponding vectors for other regions. Notably, this measures shared changes in regions’ representational *strength*, rather than through representational space (which can influence other metrics; Anzellotti & Coutanche, 2018). In this sense, the technique is content-invariant. The new method is termed here “Representational Connectivity” (RC). This term has been previously applied in reference to the degree of correlation between RDMs from multiple brain regions (Kriegeskorte *et al*., 2008b). However, we believe it is better applied in this case: “connectivity” implies a dynamic pattern of synchronized brain activity, which may only be determined by considering change over time. In contrast, this second-order RSA collapses the time domain, and thus is not sufficient to infer network activity on its own. Our RC analysis adds a time dimension to RSA that allows us to test the extent to which regions are subject to the same fluctuations over time. In this study, we ask how *representational networks* identified using this method overlap and are distinct from the representationally similar regions that are identified via second-order RSA, or with functionally connected regions identified via a univariate FC analysis.

## 2. Materials and Methods

### 2.1 Participants and Procedure

We applied the novel RC analysis method to existing data from Nastase et al. (2017), made available on OpenNeuro (https://openneuro.org; Poldrack and Gorgolewski, 2017). This study collected fMRI data from twelve right-handed adults (5 males, mean age = 25.4, standard deviation = 2.6).

Participants viewed video clips of five different animal taxonomies (birds, ungulates, primates, insects, and reptiles) performing each of four behaviors (fighting, swimming, eating, and running) for a total of 20 stimulus conditions (Figure 1). Each condition had two unique clips and all clips were flipped horizontally to create a total of 80 stimuli. For each of 10 runs in the experiment, participants were instructed to focus on either the behavior (five runs) or the taxonomy (five runs) in each clip and indicate via button press when a taxonomy or behavior was repeated. Run order was counterbalanced across participants. In a condition-rich ungrouped-events design, each trial was presented as one silent 2-second video clip followed by a 2-second fixation period. Each run began and concluded with a 12-second fixation cross, and included four behavior repetition events and four taxonomy repetition events for a total of 88 experimental trials (Figure 2).

**Figure 1.**
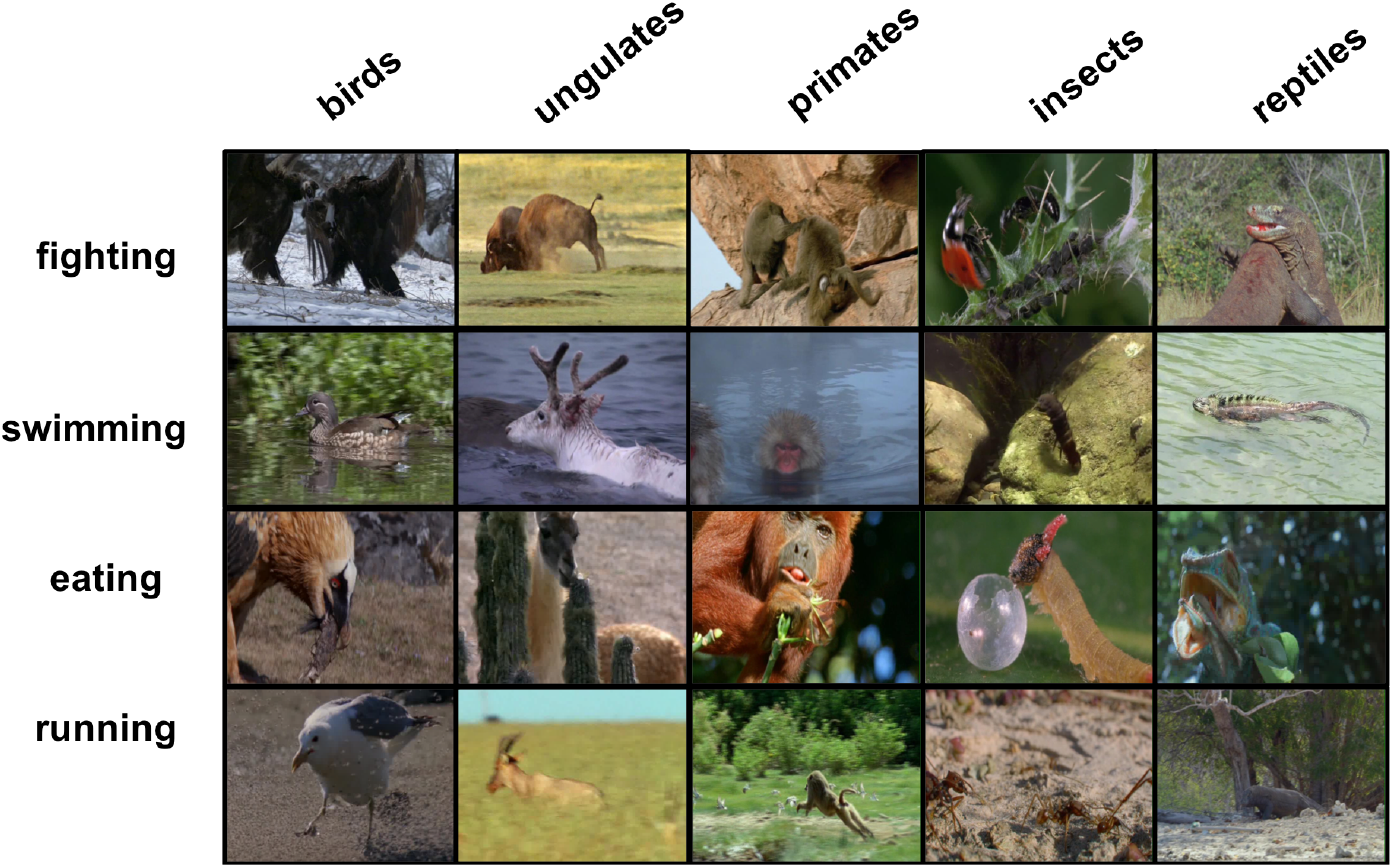
Presented videos featured 4 behavior and 5 taxonomy conditions.

**Figure 2.**
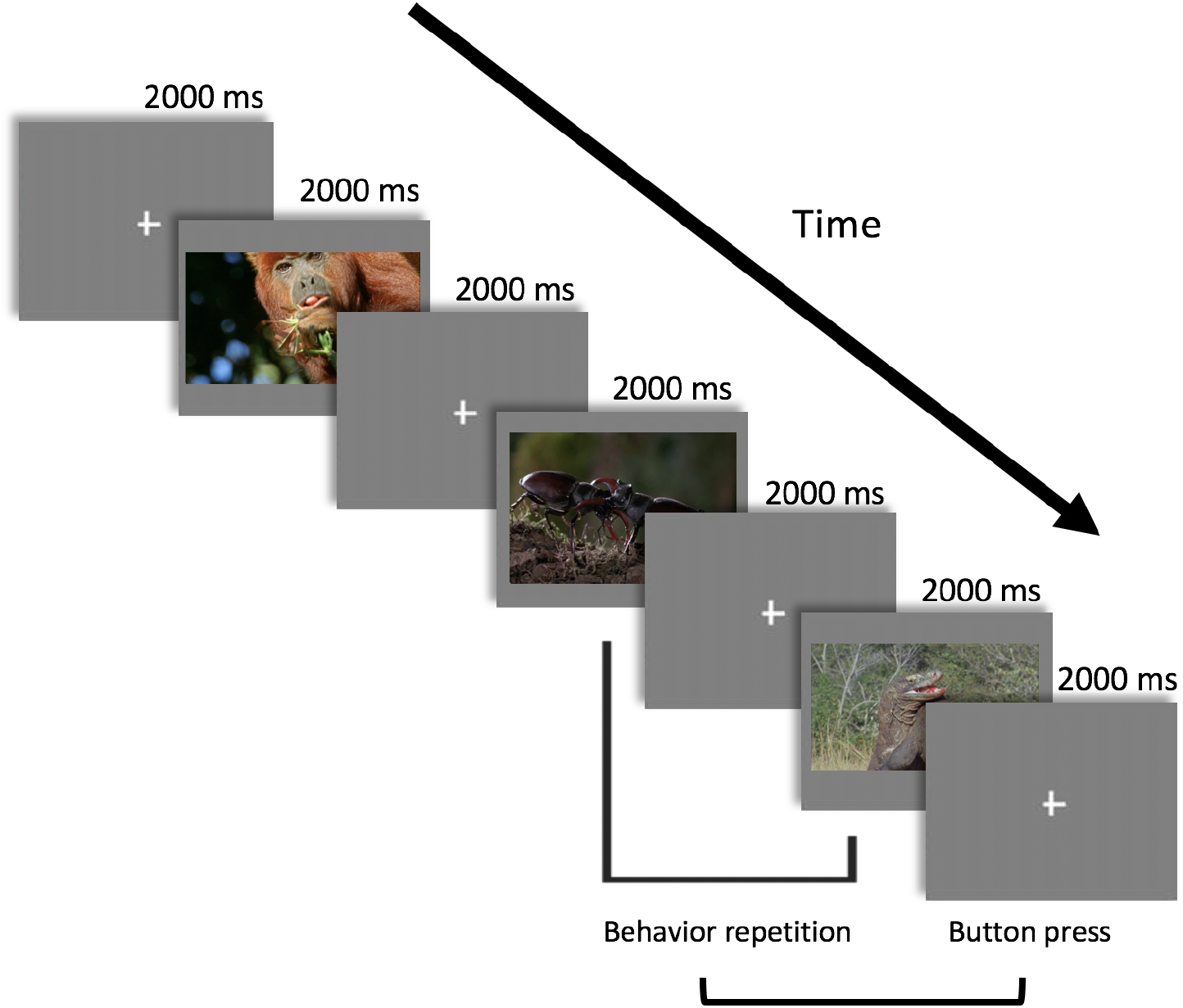
Experimental design. Here, the participant would be completing the behavior task.

### 2.2 Imaging Acquisition and Preprocessing

Structural and functional images were collected with a 3T Philips Intera Achieva MRI scanner. Blood-oxygenation-level-dependent (BOLD) response levels were collected through gradient echo-planar images (TR = 2000, TE = 35ms, resolution = 3mm^3^ isotropic, flip angle = 90^0^, and matrix size = 80 x 80) in an interleaved acquisition order.

Preprocessing was conducted in Freesurfer (Fischl et al., 2002; Fischl et al., 2004) and AFNI (Analysis of Functional NeuroImages) (Cox, 1996), and was kept as close as possible to the original parameters (Nastase et al., 2017) within the confines of our desired analyses.

We conducted cortical reconstruction and gray matter parcellation on each subject’s data using Freesurfer. Using AFNI, we applied a bandpass filter to functional images that removed frequencies below 0.0067 Hz and above 0.1 Hz. To analyze functional time series data on a trial-by-trial basis, we used AFNI’s 3dDeconvolve to calculate beta coefficients for all TRs during which stimulus clips were presented (in this case, one TR per trial). VT was selected as a seed region. VT is implicated in the representation of higher-level visual and conceptual features within multi-voxel patterns (Coutanche, Solomon, & Thompson-Schill, 2016; Haxby et al., 2001), including taxonomy- and species-relevant information for animal stimuli (Connoly *et al*., 2012; Coutanche and Koch, 2018). We defined VT within individuals based on a procedure outlined in Coutanche and Koch (2018) in which the region is defined as extending from 20 to 70 mm posterior to the anterior commissure, incorporating the lingual, fusiform, parahippocampal and inferior temporal gyri. We defined white matter by extracting the left/right white matter regions from the Harvard-Oxford subcortical structural atlases (Frazier et al., 2005). Gray matter was defined by referencing the automated Freesurfer segmentation atlas. All anatomical and functional images for each participant were then standardized from native space to MNI152-space using AFNI.

### 2.3 Representational Connectivity Analysis

Further analyses were conducted in MATLAB with the Princeton MVPA toolbox (Detre *et al*., 2006) and custom scripts. Using the beta coefficients for each voxel derived from AFNI, we estimated an activity pattern for each of the 20 stimulus clip conditions within each individual. To calculate an RDM, a leave-one-run-out cross-validation framework was used to hold out the patterns from each run and compare them with averaged pattern data from the other four runs in that attentional condition (behavior or taxonomy). As a metric of similarity, we calculated the correlation between the pattern data for each stimulus clip condition and the patterns for each of the 20 stimulus clip conditions The results were organized into a 20×20 RDM for each run by averaging the RDMs from each iteration to create a whole-series RDM with each cell representing the correlation between pairs of stimulus clip conditions.

To quantify changes to representational similarity strength over time, we first quantified the influence of removing each trial of the whole-series RDM for each run on the activity patterns in the VT seed region. For this comparison, we applied jackknife resampling to data from within VT. The jackknife (JK) technique is accomplished via the following procedure, which was implemented separately for each run:

1. One trial is removed from the functional data. A new RDM is then created from the remaining data as above.
2. The new trial-removed RDM is correlated with the original whole-series RDM for the run.
3. These steps are repeated for each trial. The resulting correlation coefficients are concatenated into a vector representing the influence of each trial on the whole-series RDM for that run.

After repeating this procedure for all trials in each run (88 here), the resulting modulation vectors are then concatenated for all runs in each condition (here, a vector of 440 values for five runs, for a region) (Figure 3a). In this study, we conducted this procedure for each attentional condition (behavior and taxonomy) separately, giving us JK vectors that capture the influence of each trial within VT for each task.

**Figure 3.**
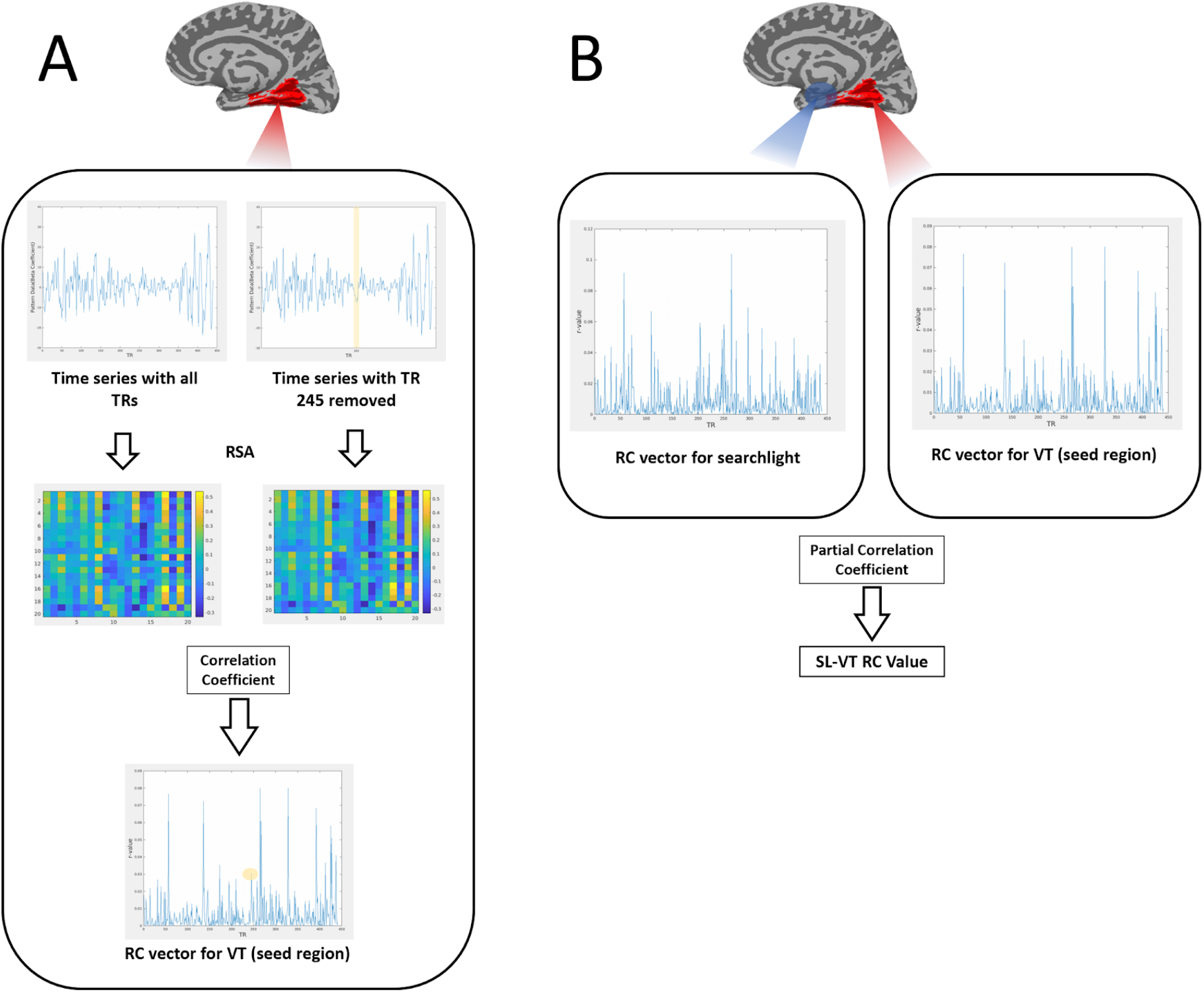
Flow diagram of Representational Connectivity Analysis. (A) After generating the whole-series RDM incorporating all TRs in a run for the seed region (VT), RC RDM (consisting of an RDM with a specific TR removed) is generated for each TR. RC RDM is then correlated with the original whole-series RDM for the run. The correlation coefficients representing the modulation of every TR that was sequentially removed are collated into a single vector. (B) This process is repeated for each searchlight, and the resulting vector is correlated with the VT vector to calculate that searchlight’s representational connectivity value.

To analyze the relationship between activity throughout the brain and activity within VT, a whole-brain 3-voxel-radius searchlight (Kriegeskorte, Goebel, & Bandettini, 2006) was created for gray matter voxels. For every searchlight, the JK resampling procedure applied in VT was repeated. Pattern data within each searchlight was reorganized in the same 20×20 RDM and then JK resampled to obtain a 440×1 modulation vector. As we aimed to quantify regions in which fluctuations in representational strength were synchronized with VT, we then correlated the modulation vector obtained for each searchlight with the modulation vector obtained for VT while partialling-out white matter. White matter was partialled-out to minimize the effect of scanner drift and non-neuronal signal change on the final connectivity result. The partial correlation coefficients obtained for each searchlight were collated into whole-brain representational connectivity (RC) maps for the behavior and taxonomy conditions within each individual (Figure 3b). After a Fisher-Z transformation of the partial-correlation whole-brain RC map for every participant, a second-level paired t-test was performed with the transformed behavior and taxonomy maps.

### 2.4 Second-order RSA

Results from the RC analysis were compared with results from a second-order RSA (Kriegeskort et al., 2008) of the same data that did not include jackknife resampling. Using the leave-one-run-out cross-validation framework for the behavior and taxonomy attentional conditions, a 20×20 RDM of all stimulus clip conditions was calculated for voxels within the VT ROI and each searchlight. The resulting RDMS for behavior and taxonomy within VT were correlated with RDMs for their respective conditions within each searchlight. White matter was partialled-out in our second-order RSA in the interest of keeping results for the RC and RSA methods as consistent as possible. The partial correlations obtained were collated into wholebrain maps of representational similarity for behavior and taxonomy for each individual. Following a Fisher-Z transformation of individual partial-correlation RSA maps, we generated group maps for each condition with significance thresholds of p<0.001 and p<0.005. To assess task-related differences in overall representational similarity, the analysis concluded with a second-level paired t-test of behavior vs. taxonomy maps at each significance threshold.

### 2.5 Functional Connectivity

RC results were also compared to results from a typical FC analysis (Fair et al., 2006) of the same data. Using AFNI, we extracted the mean BOLD activation time-series for our TRs of interest (TRs during which stimulus clips were presented) within the VT ROI. We then conducted a whole-brain searchlight partial correlation of each attentional condition, using the VT time-series and searchlight time series as white matter signal is partialed out. The results of the analysis represent the correlation of each voxel’s activation time-series with the mean timeseries of VT, and a significant correlation indicates BOLD signal synchronization between voxels and VT. Following a Fisher-Z transformation of the individual FC maps, group FC maps for behavior and taxonomy were calculated with significance thresholds of p<0.001 and p<0.005 to give a cluster-corrected threshold of p<0.05. The FC analysis also concluded with a second-level paired t-test of behavior vs. taxonomy at each threshold.

## 3. Results

Using the group-level processed data, we created three sets of whole-brain maps with VT as a seed region. We applied the novel RC analysis method to identify representational networks of regions whose fluctuations in representational similarity were linked to VT; we conducted a conventional RSA in which all timepoints were collapsed for a global RDM to identify regions with similar overall representational spaces to VT; and we conducted a classic FC analysis to identify regions whose fluctuations in BOLD signal were synced with VT. For each method, we created cluster-corrected maps for each attentional condition (behavior or taxonomy) at a strict significance threshold of p<.001 and a more liberal threshold of p<.005. We then assessed the degree of overlap between the representational network and each of the other networks for each attentional condition at each threshold. Finally, we assessed possible task-related differences in each map via a second-level paired t-test of behavior vs. taxonomy runs.

### 3.1 Representational Connectivity

The RC method successfully identified a network of brain regions that exhibited fluctuations in representational similarity shared with fluctuations in VT. Figure 4 shows the regions in which the JK modulation vector was significantly correlated with the modulation vector for VT. At a conservative threshold of p<0.001, behavior and taxonomy maps both show a network of significant RC within lower-level primary and secondary visual regions, including the occipital pole and area of extrastriate cortex in both hemispheres. Significant RC is also observed in motor and somatosensory regions of dorsal frontal/parietal cortex. The network also includes some higher-level cognitive regions, including the right superior frontal gyrus, and posterior parts of the intraparietal sulci. Considering a more liberal threshold of p<0.005, we observe a more pronounced effect in dorsal stream regions and superior structures, including left superior parietal cortex, left superior temporal sulcus, left insula, and right posterior cingulate cortex. In Table 1, we report the RC method’s peak coordinates for clusters in the taxonomy attentional condition. These coordinates were not significantly different for the behavior condition. The overlap of these coordinates with the FC and the second-order global RSA group maps are also reported.

**Figure 4.**
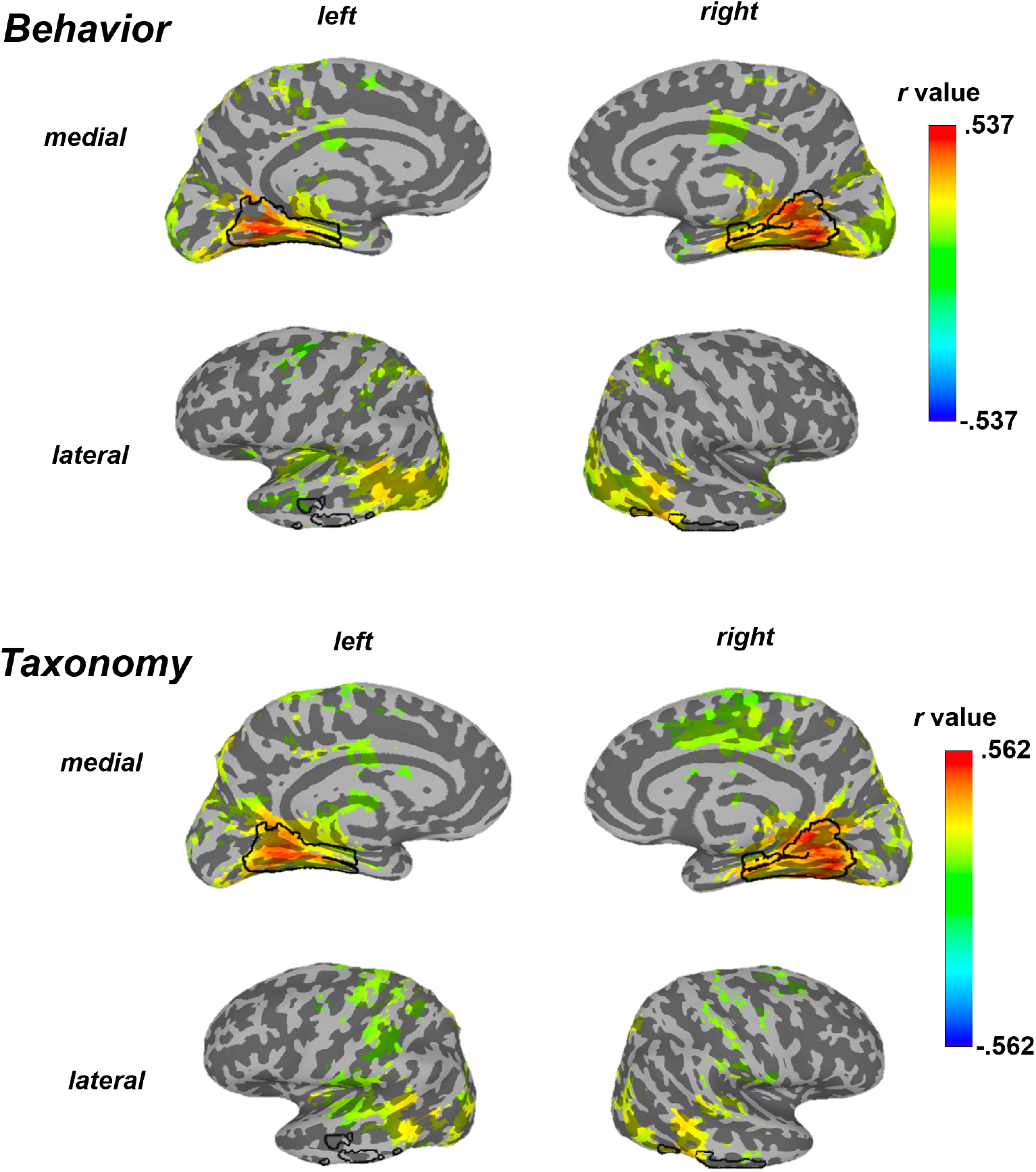
Surface maps showing regions with significant RC to VT for the Behavior-focused (top) and Taxonomy-focused (bottom) conditions. The VT seed region is outlined on each surface.

**Table 1.** Peak coordinates of Taxonomy-focused clusters with averaged correlation coefficient values and voxel size from the jackknife Representational Connectivity (RC) method, Functional Connectivity (FC) and second-order global RSA. Coordinates of peak correlation in each cluster and sub-clusters are reported. All clusters significant at 0.005; * = significant at 0.001. Peak coordinates within the VT seed region are not included in the table.

|  | Region | Coordinates (MNI152) |  |  | Volume (voxels) | RC corr. coeff. | Overlap |  | alpha |
| --- | --- | --- | --- | --- | --- | --- | --- | --- | --- |
|  |  | X | Y | Z |  |  | RSA | FC |  |
| **Cluster Peak:** | **Right posterior lingual gyrus** | **16** | **-62** | **-5** | **27,061** |  | **yes** | **yes** | **<<0.01** |
| Sub-peaks: | Left superior temporal sulcus | -59 | -65 | 7 |  | 0.352 | yes | yes |  |
|  | Left intraparietal sulcus | -17 | -63 | 63 |  | 0.342* | no | yes |  |
|  | Left extrastriate cortex | -48 | -74 | 16 |  | 0.34* | yes | yes |  |
|  | Right cingulate gyrus | 14 | -43 | 61 |  | 0.325* | no | yes |  |
|  | Left angular gyrus | -30 | -76 | 44 |  | 0.318* | yes | yes |  |
|  | Left paracentral lobule | -10 | -36 | 75 |  | 0.316 | no | yes |  |
|  | Right extrastriate cortex | 41 | -76 | 10 |  | 0.309* | yes | yes |  |
|  | Right intraparietal sulcus | 22 | -62 | 58 |  | 0.292* | yes | yes |  |
|  | Left Insula | -42 | -8 | -5 |  | 0.287 | no | no |  |
|  | Right dorsal posterior cingulate cortex | 2 | -31 | 38 |  | 0.256 | no | no |  |
|  | Occipital pole | 3 | -95 | 10 |  | 0.233* | yes | yes |  |
|  | Right precentral gyrus | 22 | -18 | 76 |  | 0.213* | no | yes |  |
|  | Left postcentral gyrus | -47 | -38 | 65 |  | 0.256 | no | yes |  |
|  | Right supramarginal gyrus | 60 | -21 | 40 |  | 0.198* | no | yes |  |
|  | Right superior frontal gyrus | 28 | 16 | 63 |  | 0.166* | no | no |  |

|  | Region | Coordinates (MNI152) |  |  | Volume (voxels) | RSA corr. coeff. | Overlap |  | alpha |
| --- | --- | --- | --- | --- | --- | --- | --- | --- | --- |
|  |  | X | Y | Z |  |  | RC | FC |  |
| **Cluster Peak:** | **Right lingual gyrus** | **16** | **-62** | **-5** | **14,838** |  | **yes** | **yes** | **<<0.01** |
| Sub-peaks: | Left middle temporal gyrus | -59 | -56 | 10 |  | 0.335* | yes | yes |  |
|  | Right middle temporal gyrus | 55 | -50 | 9 |  | 0.345* | yes | yes |  |
|  | Left superior temporal sulcus | -41 | -35 | 13 |  | 0.193* | yes | yes |  |
|  | Left inferior occipital gyrus | -46 | -79 | -11 |  | 0.265* | yes | yes |  |
|  | Left angular gyrus | -30 | -81 | 43 |  | 0.287* | yes | yes |  |
|  | Left middle occipital gyrus | -40 | -89 | 12 |  | 0.234 | yes | yes |  |
|  | Right superior parietal lobule | 15 | -62 | 58 |  | 0.259 | yes | yes |  |
|  | Right precuneus | 23 | -61 | 19 |  | 0.271* | yes | yes |  |
|  | Left posterior cingulate cortex | -2 | -56 | 16 |  | 0.239* | yes | yes |  |
|  | Right cuneus | 5 | -95 | 20 |  | 0.204 | yes | yes |  |
| **Cluster Peak:** | **Right pallidum** | **19** | **-2** | **7** | **4,305** | **-0.267\*** | **no** | **no** | **<<0.01** |
| Sub-Peaks: | Left dorsal striatum | -10 | 4 | -2 |  | -0.235* | no | no |  |
|  | Right orbitofrontal cortex | 22 | 58 | -11 |  | -0.232* | no | no |  |
|  | Left orbitofrontal cortex | -27 | 62 | -16 |  | -0.186* | no | no |  |

|  | Region | Coordinates (MNI152) |  |  | Volume (voxels) | FC corr. coeff. | Overlap |  | alpha |
| --- | --- | --- | --- | --- | --- | --- | --- | --- | --- |
|  |  | X | Y | Z |  |  | RC | RSA |  |
| **Cluster Peak:** | **Right inferior fusiform gyrus** | **46** | **-53** | **-17** | **26,061** |  | **yes** | **yes** | **<<0.01** |
| Sub-peaks: | Right extrastriate cortex | -34 | -84 | 16 |  | 0.712* | yes | yes |  |
|  | Left extrastriate cortex | 37 | -79 | 18 |  | 0.684* | yes | yes |  |
|  | Right middle temporal gyrus | 52 | -68 | 11 |  | 0.642* | yes | yes |  |
|  | Right intraparietal sulcus | 33 | -60 | 58 |  | 0.577* | no | no |  |
|  | Right superior parietal lobule | 33 | -59 | 59 |  | 0.577* | no | no |  |
|  | Left superior parietal lobule | -28 | -57 | 61 |  | 0.556* | yes | no |  |
|  | Left anterior fusiform gyrus | -30 | -3 | -37 |  | 0.352* | no | no |  |
|  | Left superior frontal gyrus | -43 | -5 | 59 |  | 0.344* | yes | no |  |
|  | Right inferior temporal gyrus | 32 | 3 | -42 |  | 0.331* | no | no |  |
|  | Left inferior frontal gyrus | -45 | 11 | 31 |  | 0.315* | no | no |  |
|  | Left superior temporal gyrus | -60 | -11 | -3 |  | 0.279* | yes | no |  |
|  | Left inferior frontal gyrus | -50 | 40 | 10 |  | 0.275* | no | no |  |
|  | Right postcentral gyrus | 59 | -19 | 47 |  | 0.272* | yes | no |  |
|  | Right superior temporal gyrus | 46 | -38 | 13 |  | 0.246* | yes | yes |  |
|  | Left cingulate | -12 | -17 | 43 |  | 0.228* | yes | no |  |
| **Cluster Peak:** | **Right superior frontal gyrus** | **22** | **46** | **19** | **959** | **-0.325\*** | **no** | **yes** | **<0.02** |
| Sub-Peaks: | Right orbitofrontal cortex | 20 | 56 | -3 |  | -0.242* | no | yes |  |
|  | Left anterior cingulate | -11 | 46 | 7 |  | -0.223* | no | no |  |

### 3.2 Representational Similarity Analysis

A traditional second-order RSA, collapsing all time points across the experiment, identifies a set of midbrain temporal and occipital regions in both hemispheres with significantly similar overall representational spaces to VT. High levels of correlation between the VT RDM is observed in lower-level visual areas, including lateral occipital cortex and the right lingual gyrus, and higher-level ventral stream visual regions, including the middle temporal gyri. Similar representational spaces are also observed in cortex associated with spatial perception and multisensory integration, including cingulate cortex, the right superior parietal lobule, and left angular gyrus. Again, maps for taxonomy and behavior focus are highly similar, although some frontal activation is observed for behavior at the more liberal threshold not observed for taxonomy; again, the difference does not rise to significance.

### 3.3 Functional Connectivity

The traditional FC analysis identifies a broad network of regions associated with VT via fluctuations in BOLD activity. A wide array of functionally connected regions are observed in the primary and secondary visual cortex and in the ventral and dorsal visual streams in both hemispheres, as well as prefrontal cortex, the superior parietal lobule, and cingulate cortex. A high degree of connectivity is observed in all maps, and even at a more stringent threshold, FC reveals a pervasive functional network throughout the brain.

### 3.4 Overlap

Both RSA and FC maps show similar cluster patterns along the ventral stream and within the occipital pole where there is a high degree of overlap with the representational network (Figure 5, Figure 6).

**Figure 5.**
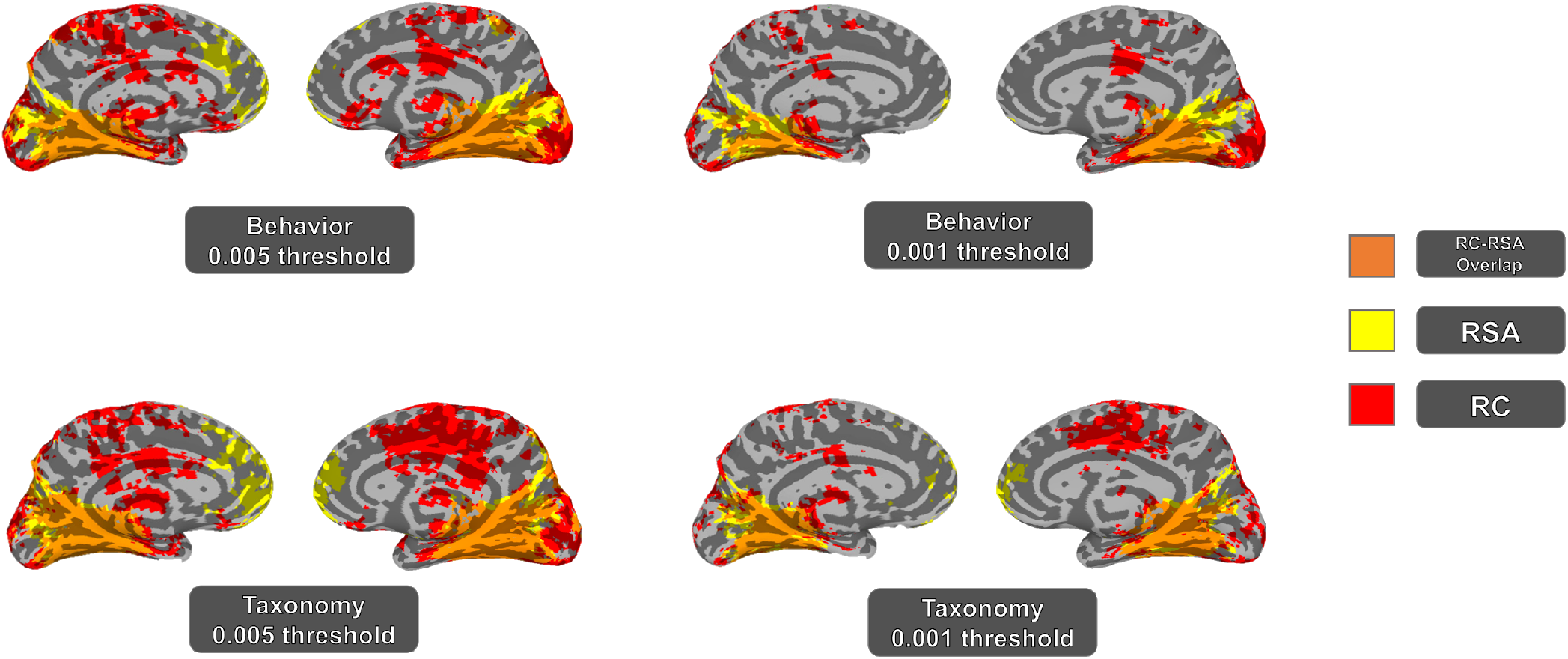
Overlap in regions associated with ventral temporal cortex through Representational Connectivity (RC) analysis and second-order representational similarity analysis (RSA) in a second-level *t*-test, separated based on attentional condition (behavior or taxonomy focus) that reached significance at p=0.001 and p=0.005.

**Figure 6.**
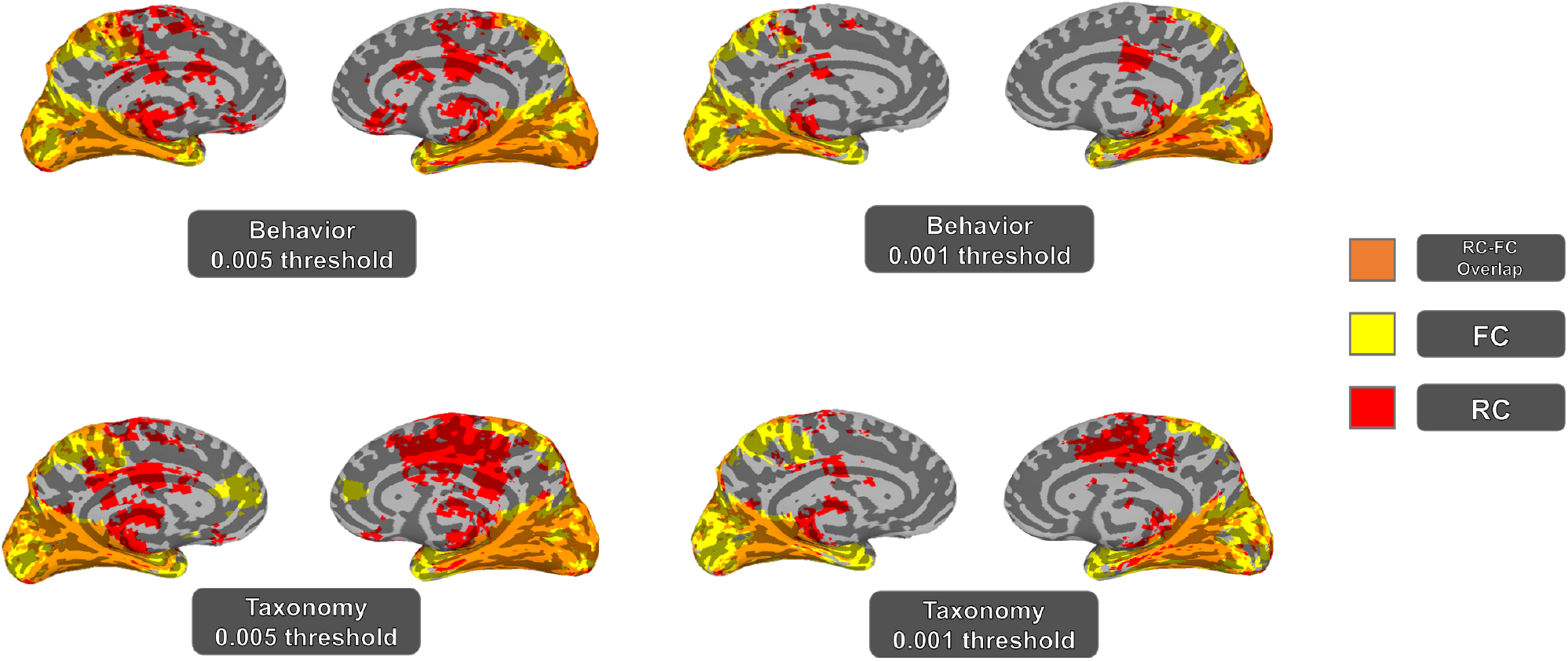
Overlap in regions associated with ventral temporal cortex through Representational Connectivity (RC) analysi and functional connectivity (FC) analysis in a second-level t-test, separated based on attentional condition (behavior or taxonomy focus), that reached significance at p=0.001 and p=0.005.

A great degree of overlap between RC and second-order RSA maps was observed in occipital regions. However, there were clusters along the dorsal temporal lobe and occipital regions of the ventral visual stream in which there was no overlap between the RC and second-order RSA maps. Some superior regions are included in the RC maps but not the RSA maps, especially at the more liberal threshold; these include higher-level frontal areas as well as motor and somatosensory cortex.

We saw few regions in the RC network that did not overlap with the FC network to some degree. This is attributable in part to the wide degree of functional connectivity observed in every map. We did observe small clusters in the middle temporal region that did not completely overlap with FC, especially when considering the more stringent threshold. We also observed small occipital clusters that were identified in the RC network but not the FC network at the more stringent threshold.There was also an overlap between the RC and FC maps in parietal regions at a threshold of p<.005; at the more stringent threshold, overlap between RC and FC networks is restricted to the ventral stream.

### 3.5 Task Conditions

Our tests did not show a significant difference between behavior and taxonomy maps at a p-threshold of either 0.001 or 0.005. As noted above, slight differences between tasks could be observed when considering RSA, FC, and RC network maps; however, our second-level paired t-test of behavior vs. taxonomy showed that this difference did not rise to significance for any of the three methods considered (p>.005). In addition, the degree of overlap between RC and RSA maps or between RC and FC maps did not differ significantly by task.

## 4. Discussion

Here, we present results from a novel method for identifying regions that are linked via fluctuations in the strength of representational similarity over time. We apply Representational Connectivity analysis to a dataset using VT as a seed region to identify brain regions with synchronized fluctuations of strength in representational similarity. Previous evidence has demonstrated that analyzing activity over time adds information above and beyond what is available from analysis of fMRI activity collapsed across the time domain (Coutanche & Thompson-Schill, 2013; Friston, 1997). We demonstrate that, as in the case of FC for univariate analysis and IC for MVPA, incorporating a time-domain analysis via RC adds information above and beyond that available in second-order RSA. Our findings illustrate the potential benefits to future RSA-based experimental studies if investigators choose to incorporate RC into analyses.

RC analysis successfully identified a network of regions linked to our seed region (VT) by shared fluctuations in representational similarity. In response to the dynamic animal behavior stimuli used in this experiment (Nastase et al, 2017), we observe a network incorporating regions associated with lower-level visual processing, spatial cognition, perceptual-motor integration, and visual attention.

The regions considered to compose VT are implicated in the representation of information for visual categories within multi-voxel patterns, including faces, animals, tools, and places, among others (Haxby *et al*., 2001, O’Toole *et al*., 2005). This experimental task demanded that subjects allocate visual attention, access category information, and hold relevant information in mind between trials. In the RC network, we observe synchronized change across areas of visual cortex from trial to trial; this relationship makes intuitive sense given that individual trials present new visual information that must be processed in order to make judgments about the stimuli. Synchronized fluctuations were also observed in the frontal cortex, anterior parietal cortex, and cingulate cortex. These regions might be involved in maintaining and allocating visual attention effectively, and managing information held in memory while focusing on the present task. Interestingly, despite this task’s primary focus on visual category representation, the RC analysis also identified regions linked to motor activity and spatial cognition: some synchronized fluctuations were observed in motor and somatosensory cortex, especially at the more liberal threshold. We also identified dorsal regions linked to VT via fluctuations in representational similarity - notably, significant RC was observed in both the left and right intraparietal sulci (IPS), which guide the integration of perceptual and motor information (Grefkes & Fink, 2005). In activation-based studies using static visual stimuli, the IPS has been shown to respond more to images of tools and manipulable objects than to images of animals (Mruczek et al., 2013). However, in this task, representational similarity in these regions shared modulation with VT for trial-to-trial changes in representational strength. RC observed here illustrates that this method may be used to identify regions in which the impact of task-relevant visual information on representational strength is linked to the impact of task-relevant information from other domains or sensory modalities. For instance, this might reflect information being passed to these regions in order to detect potential future motor actions. Alternatively, this might reflect the cognitive process required for the task, where participants must respond based on whether a behavior or taxonomy repeats in the observed videos.

When comparing the RC network to a map of regions obtained from a second-order RSA that collapsed data across all timepoints, we found partial overlap between RC and RSA but also a number of regions, including frontoparietal areas, that were unique to the RC network. These regions were linked to VT via shared fluctuations in representational similarity but did not have significantly similar representational spaces. The nature and function of this association remains to be investigated, but it is clear that adding the time domain as an element of analysis has contributed information above and beyond what was available from RSA alone. This also indicates that it is possible to detect common fluctuations, even if these spaces are governed by unique dimensions (e.g., if a particular time-point marks a dramatic shift, this can be observed across spaces).

The second-order RSA also revealed some regions with a shared representational space, but without shared across-time fluctuations. This included parts of the dorsal temporal lobe and regions of orbitofrontal and prefrontal cortex. Regions with synchronized fluctuations identified via RC were affected in the same manner as VT by the removal of specific trials, *i.e*., their RDMs were modulated to the same degree in response to the same stimuli. If two regions are shown via second-order RSA to have dissimilar representational spaces, they may still be correlated in terms of the degree to which those spaces are affected by specific items; likewise, if two regions are not synchronized in terms of their change in response to specific items, they may still have highly similar representations for those items overall.

When comparing the RC network to a network obtained via FC analysis of regions whose fluctuations in BOLD signal were synchronized over time, we found a larger degree of overlap than with the second-order RSA, as most regions in the RC network were also linked to VT in the FC analysis. However, we identified some clusters that were linked to VT in the RC network but not in the functional network. These regions’ BOLD signals are not significantly synchronized with VT over time, but they do share fluctuations in representational similarity. Within regions linked by FC, which are presumed to have related responses and functional roles during this task, there are smaller regions also linked by their fluctuations in representational similarity. Additional regions may not be functionally linked but are associated in RC analysis, indicating a possible information-sharing mechanism unique to these regions that may or may not be reflected by univariate functional connectivity. Some previous findings show that even when task-dependent univariate signal falls to baseline, multi-voxel pattern information still reveals effects of task in the same cortical regions. For instance, Harrison and Tong (2009) illustrated that visual stimuli held in mind could be decoded accurately from multi-voxel pattern information in V1 – V4 even after a long delay during which overall BOLD activity fell to baseline levels. If such patterns may be observed for overall task-based classification, it is highly probable that the same holds for FC versus RC. Similarly, comparisons between IC and FC reveal regions that are connected through shared fluctuations in multi-voxel patterns, without having associated FC (Coutanche and Thompson-Schill, 2013; 2014).

We did not find a significant difference between attentional conditions (behavior vs. taxonomy) at a cluster-corrected threshold of p<.005. Based on the nature of the behavior/taxonomy task itself, there was an assumption that the conditions would show some degree of difference. On the one hand, this agreement is a helpful and positive signal of reliability for the jackknife approach, given that these are independent runs (*i.e*., within-subject replication). On the other hand, this may indicate that this task is not modulating the shared networks. It is possible that our seed (VT) was not sufficient to detect a task-related difference; RC networks defined using a different seed region may capture the task-related factors that modulate signal in this case. The original authors report different regions of peak mean searchlight accuracy and overall differences in classification accuracy between attentional conditions at the group level, but they did not perform a direct test between behavior and taxonomy data and noted similarity in the relevant networks (Nastase et al, 2017).

Based on the results presented here, in combination with past evidence of the utility of connectivity analyses for univariate and multivariate studies, we propose that RC could be considered in future RSA-based studies to add a time dimension, in a similar way as for univariate analyses (FC) and MVPA (IC). Incorporating RC identifies new connections because regions associated solely based on having a similar representational space (i.e., second-order RSA) will not necessarily share synchronized changes in representational strength over time, and vice versa. Similarly, although other methods have examined whether regions share changes through representational space (Anzellotti & Coutanche, 2018), this new method examines shared changes in representational *strength*, rather than content, allowing it to identify RC between regions that have distinct principles underlying representational similarity. Overall, this method fills a gap in the current set of fMRI analysis tools, and represents a new lens through which to view brain activity on a network level.

## Acknowledgments

We thank Mac Shine for invaluable discussions during the conception of this project, and thank Heather Bruett, Griffin Koch, John Paulus and Xueying Ren for conversations related to the work. We are also grateful to Samuel Nastase and co-authors for making their dataset available.

